# TWEAK is increased in ulcerative colitis and contributes to fibroblast-mediated monocyte activation via heterologous non-canonical NF-kB/STAT3 signalling

**DOI:** 10.1101/2025.09.23.678006

**Authors:** Carlos Matellan, Bella Raphael, Gareth R. Jones, Mary Nwaeziegwe, Sarah Balfe, Cian M Ohlendieck, Cristina Bauset, Méabh B Ní Chathail, Ciarán Kennedy, Katie Doogan, Des Winter, Carol Aherne, Helen M Roche, Calum Bain, Stephen D. Thorpe, Glen Doherty, Mario C. Manresa

## Abstract

**Background and Aims:** Interactions between fibroblasts and monocytes have emerged as a contributing factor in IBD pathogenesis and therapy resistance, owing to the ability of both cell types to participate in tissue inflammation and repair. We have previously shown that the TNF superfamily factor TWEAK (TNFSF12) can induce a UC-like inflammatory profile in colonic fibroblasts *in vitro*, in turn promoting monocyte adhesion and activation. However, the mechanisms underlying fibroblast-monocyte communication and its dysregulation in colitis are incompletely understood.

**Methods:** Here we use co-culture models, human biopsies from ulcerative colitis (UC) patients and healthy donors, and public single-cell transcriptomics to characterise the mechanisms underlying fibroblast-mediated monocyte activation.

**Results:** We show that TWEAK-treated inflammatory fibroblasts induce a transcriptional programme that resembles early monocyte/macrophage intermediates in UC and is enriched for genes associated with resistance to anti-TNF (*TREM1, OSM, IL1B*) and susceptibility to IBD *(NOD2, ATG16L1*). We find that conditioned media from TWEAK-treated fibroblast causes a sustained activation of STAT3 phosphorylation in monocytes, and that inhibition of the NF-κB Inducing Kinase (NIK) impairs the ability of inflammatory fibroblasts to activate STAT3 phosphorylation in monocytes, resulting in reduced expression of inflammatory mediators. Using tissues from UC patients, we show that the expansion of CD90^+^/PDPN^+^ inflammatory fibroblasts in UC correlates with the accumulation of TWEAK^+^ myeloid cells in the colonic mucosa, and that these fibroblasts co-localise with infiltrating monocytes in sites of active inflammation.

**Conclusion:** Together, our findings suggest that the TWEAK/NF-κB/STAT3 axis represents an attractive target to tune inflammatory stroma/monocyte crosstalk.

## INTRODUCTION

Inflammatory bowel diseases (IBD) are chronic inflammatory disorders of the gastrointestinal tract, principally comprising ulcerative colitis (UC) and Crohn’s disease (CD). IBD is a global healthcare challenge, currently affecting ∼15 million people globally^1^ and causing the loss of over 1.5 million disability-adjusted life-years (DALYs)^2^, with rising incidence in both western and newly industrialised countries.

Since the approval of the first biological therapies targeting TNF in IBD, a variety of different treatments have become available including anti-α4β7 integrins, anti-IL-12/IL-23 and JAK/STAT inhibitors, with several others undergoing clinical trials^3,4^. Yet, despite the diversity of options, these treatments still present multiple limitations. First, a significant proportion of patients (10-40% for anti-TNF) are initially non-responsive to treatment for reasons that are poorly understood (primary non-response)^5^. Second, it is common for primary responders to develop secondary loss-of-response (∼20% per year), becoming refractory over time even when the treatment initially succeeds at achieving remission^5,6^. Moreover, available drugs focus on managing the inflammatory response but often fail to address mucosal healing, and none of the currently available therapies specifically target resident structural cells, which is likely critical to improve mucosal healing and long-term remission^7^. Indeed, understanding the mechanisms whereby structural cells contribute to IBD pathogenesis could provide new therapeutic avenues to overcome current limitations.

While the pathogenesis of IBD is thought to be complex and multi-factorial, with no cell type being solely responsible, a growing body of evidence points towards monocytes/macrophages as an interesting target due to their dual role in gut inflammation and tissue repair. Indeed, tissue resident macrophages carry out tolerogenic functions in the gut and are steadily replenished by bone-marrow derived circulating monocytes^7^. During inflammation, however, resident homeostatic macrophages are rapidly replaced by inflammatory macrophages (CD14^hi^) that are generated from the novo recruited circulating monocytes and accumulate in the mucosa of IBD patients^8,9^. These secrete pro-inflammatory cytokines such as IL-1β, IL-6, TNF or IL-23^10,11^. Interestingly, many IBD susceptibility loci are associated with macrophages (*NOD2, PTPN2, TNFAIP3, STAT3*)^12^, and several macrophage markers/products (*TREM-1, OSM*) are associated with therapy resistance, pointing towards macrophage dysfunction’s central role in IBD. Conversely, macrophages are also critical during inflammation resolution, contributing to the clearance of debris and commensal bacteria and secreting pro-resolving mediators that assist in epithelial regeneration and wound healing. Animal studies have shown that depletion of macrophages (*csf1*^op/op^)^13^ or dysregulation of their function (CD68TGF-βDNRII)^14^ can impair mucosal healing. Likewise, therapeutic success in IBD patients appears to correlate with decreased inflammatory macrophage infiltration and increased pro-resolution activity^15^. However, the mechanisms driving the sustained monocyte recruitment and the shift from homeostatic and restorative macrophages to a pathogenic phenotype in IBD represents an additional current gap in the knowledge.

In recent years, the intestinal stroma has emerged as a key player in IBD. As part of this newly appreciated remodelling of the stroma, a population of inflammatory fibroblast identified by markers such as Podoplanin (PDPN)^16,17^, and by the expression of multiple cytokines and chemokines, has been identified in several studies. These cells expand in UC^17,18^ and in CD^19^, and their emergence is associated with resistance to anti-TNF therapy in both disease endotypes^16,19,20^, highlighting their pathogenic and therapeutic relevance. Inflammatory fibroblasts may contribute to IBD by secreting cytokines and chemokines and expressing cell adhesion molecules that enable them to interact with both innate^21^ and adaptive^22^ immune cells. Indeed, studies from CD have already correlated the abundance of inflammatory fibroblasts with that of macrophages^23^. Thus, a new area of interest in the quest to improve IBD therapy is that of unearthing the key mediators and molecular mechanisms driving pathogenic fibroblast differentiation. In recent years, cytokines such as interleukin (IL)-1 or oncostatin-M (OSM) have been shown to contribute to this process through yet unknown mechanisms. In this respect, we have previously established the TNF superfamily factor TWEAK (TNFSF12) as a potential driver of inflammatory polarisation in colonic fibroblasts *in vitro*, inducing a transcriptional programme that resembles the inflammatory stroma in UC^24^. While the cellular source of TWEAK *in vivo* remains unclear^25^, we found that these inflammatory fibroblasts promote monocyte adhesion and polarisation (CD14, CXCL1) *in vitro*. Expanding on this, here we use co-culture models, human biopsies from UC patients, and available single-cell data to characterise the mechanisms underlying fibroblast-mediated monocyte activation and to assess the clinical relevance of the TWEAK-Stroma-Monocyte axis. We identify a TWEAK-dependent heterologous non-canonical NF-κB/STAT3 crosstalk pathway that drives monocyte differentiation and expression of inflammatory mediators such as *OSM*, *IL-12* or *NOD2*. Moreover, we find that fibroblast-dependent monocyte activation can be impaired by inhibiting non-canonical NF-κB signalling in fibroblasts, which emerges as a molecular target with potential for therapeutic intervention in UC.

## MATERIALS AND METHODS

### Clinical Samples

This prospective single centre study enrolled individuals with IBD undergoing colonoscopy, including diagnostic and screening colonoscopies. The study was approved by the St Vincent’s Healthcare Group, Ethics and Medical Research Committee. All participants provided written informed consent and were consecutively recruited into the study. The inclusion criteria included participants between 18-75 years old, with a known diagnosis of IBD and a select number of healthy controls with no history of IBD. Fresh colon samples were collected using standard means during colonoscopy, stored in Roswell Park Memorial Institute medium (RPMI-1640, Gibco, Waltham, MA, USA) and transported on ice for processing, or fixed in formalin and later embedded in paraffin for histology-based analyses. For patients with active disease, two biopsies were collected, one from an active region, and one from an endoscopically non-involved region. For patients in remission or healthy donors, only one biopsy was collected.

### Cell Culture and Reagents

Human primary colonic fibroblasts (CCD-18CO, American Type Culture Collection (ATCC), Manassas, VA) were cultured in Eagle’s Minimum Essential Medium (EMEM, ATCC) supplemented with 10% Foetal bovine serum (FBS, BioSciences, Ireland). Human primary colonic fibroblasts (hFBLs) were isolated from colonic resections as explained below and subsequently cultured in DMEM/F12 medium supplemented with 10% FBS, penicillin/streptomycin (100 µg/mL), gentamycin (100 µg/mL) and amphotericin B (2 µg/mL). For experiments, either CCD-18Cos or hFBLs from donors were serum deprived (in serum free EMEM or DMEM/F12 respectively) for a minimum of 6 hours, followed by stimulation with 50 ng/mL of TWEAK (PeproTech, UK) in serum-free medium for the specified time points (24 or 48 hours). For inhibition studies, CCD-18Co fibroblasts were pre-treated with NIK SMI1 (10µM, Sigma Aldrich, Cat no. SML3129) or vehicle (DMSO, Thermo Scientific, Cat no. 85190) 30 minutes before the addition of TWEAK, and then co-incubated with both stimuli for 48 hours. For Fn14 expression experiments, CCD-18Co fibroblasts were serum deprived as before and treated with 5ng/ml TNF (PeproTech, UK) for 24 hours before analysis. THP-1 cells (ATCC) were cultured in RPMI-1640 supplemented with 10% Foetal bovine serum, 1% L-glutamine (Thermo Scientific, Waltham, MA, USA) and 1% Penicillin-Streptomycin (Gibco).

### Primary fibroblast isolation

Primary human colonic fibroblasts were isolated from healthy areas of intestinal resections from colorectal cancer patients as in^26^. Briefly, colonic resections were placed in DMEM/F12 supplemented with FBS 20%, penicillin/streptomycin (100 µg/mL), gentamycin (100 µg/mL) and amphotericin B (2 µg/mL) for 30 minutes, and then washed in HBSS/EDTA/FBS (Hank balanced salt solution + 2mM EDTA + 0.5% FBS) to eliminate debris. Small pieces of the mucosal layer (5x5 mm) were sectioned from the specimens, washed vigorously in HBSS/EDTA/FBS, followed by two washes in PBS. These were placed on a petri dish that was previously scored with a needle to facilitate attachment, and cultured in DMEM/F12 supplemented with FBS 20%, penicillin/streptomycin (100 µg/mL), gentamycin (100 µg/mL) and amphotericin B (2 µg/mL) for fibroblast expansion. Fibroblast phenotype was subsequently confirmed by flow cytometry (Based on expression of CD90 and PDPN).

### Primary monocyte Isolation

Primary monocytes were purified from peripheral blood mononuclear cells (PBMC) from 5 healthy human donors following written informed consent. Whole blood was collected in tubes with EDTA (VACUETTE, Greiner, Monroe, NC, USA), and isolated using our previously reported method^24^.

### Co-culture studies

For direct co-culture experiments for bulkRNAseq, human primary colonic fibroblasts were stimulated with vehicle or TWEAK for 24 hours as previously described^24^. Next, 1/3 of the culture medium was removed and replaced with RPMI (10% FBS, 1% L-Glutamine, 1% Penicillin-Streptomycin) containing half a million THP-1. Cells were co-cultured for 24 hours, after which the THP-1 cells still in suspension were harvested in TRIzol reagent (Bio-Sciences Ltd, Ireland) for downstream RNAseq analysis.

To analyse the effects of fibroblast conditioned medium on THP-1 cells or primary monocytes, primary colonic fibroblasts were stimulated with TWEAK for 48 hours, with or without the NIK inhibitor NIK SMI1 (10 µM), as explained before, and their supernatants were stored at -80°C. Subsequently, THP-1 were resuspended in conditioned media from the fibroblasts diluted 1:1 with RPMI (with 10% FBS, 1% penicillin/streptomycin, and 1% L-glutamine) and cultured for 24h, before being lysed in either TRIzol (for RTqPCR) or radio-immunoprecipitation assay (RIPA) buffer with protease inhibitor cocktail (for Immunoblotting).

### Bulk RNAseq analysis

Samples were processed using the NextFlow nf-core/RNAseq pipeline (v3.9)^27,28^. Briefly, raw FASTQ files were trimmed using Trim Galore^29^. Trimmed reads were then mapped to human genome assembly GRCh38 and quantified at the transcriptome level using STAR^30^ and SALMON^31^. FastQC (2015)^32^ and MultiQC^33^ were used to provide QC information throughout. Downstream analysis was performed using R (v4.1)^34^. Count normalization and differential expression was performed using DeSeq2(v1.34)^35^ and gene set enrichment analysis was performed with the Enrichr web server^36^. Data was considered significant if pAdj≤0.05 and Log2fold>0.6.

### Single-Cell RNAseq analysis

Single-cell RNA sequencing data was reanalysed from Smillie *et al*^18^. Preprocessing of the data was performed using Seurat v5.2.1 according to the workflow suggested by the Satija laboratory. Raw count matrices were downloaded (https://singlecell.broadinstitute.org/single_cell) then imputed into the R environment (v3.1) and analysed using Seurat. Normalization was performed using regularized negative binomial regression via the SCTransform function, including the removal of confounding variation arising from mitochondrial mapping percentage. Doublets were identified and removed using the “Doublet Finder” package via artificial next nearest neighbour analysis. Batch effect correction was assessed using the Harmony package. Identification of highly variable genes, next nearest neighbour, clustering functions, and Uniform Manifold Approximation and Projection (UMAP) visualization were all performed using the Seurat package. Marker genes per identified subpopulation were found using the findMarker function of the Seurat pipeline.

### RT qPCR

For RNA extraction, cells were lysed on TRIzol reagent (Bio-Sciences Ltd, Ireland) and RNA was purified through a phenol-based method. RNA was reverse transcribed using the High-Capacity cDNA Reverse Transcription Kit (Life Technologies Europe BV, Netherlands) according to the manufacturer’s instructions. qRT-PCR was performed in a QuantStudio™ 5 Real-Time PCR System (Thermo Fisher Scientific) using SYBR green. The list of primers used can be found in Supplementary Table S2.

### Protein analysis

For immunoblotting analysis, whole-cells lysates were collected using RIPA buffer with protease inhibitor cocktail, and Western Blot was conducted as previously described^37^. See supplementary table S2 for a list of antibodies.

### Flow Cytometry

Colonoscopy biopsies (5x5 mm) from IBD patients or healthy donors were analysed. Biopsies were sectioned into smaller fragments and digested in 0.5ml of Milli-Q H_2_O containing 1 mg/mL collagenase type IV (C4-28, Merck) for 45 minutes at 37°C with shaking. Enzyme activity was quenched with 2 ml of FACs buffer (3% FBS and 0.5mM EDTA in PBS) before straining the resulting cell suspension through a 40-µm pore nylon membrane. Single cell suspensions were centrifuged at 2500 rpm for 10 minutes at 4°C, resuspended in FACs buffer and stained with appropriate antibody panels. Samples were stained with fluorochrome-conjugated antibodies in FACS buffer at room temperature for 30-60 minutes. For intracellular staining, surface markers were stained as above before the samples were fixed/permeabilised using the Foxp3 Transcription Factor Staining kit (Cat. No 00-5523-00, eBioscience) for 15 minutes. Fixed samples were stained with appropriate antibodies in permeabilization buffer at room temperature for 30min according to manufacturer instructions. Flow cytometry was performed with a CytoFLEX LX (Beckman Coulter), and the data analysed in FlowJo® (LLC, BD Life Sciences).

### Immunofluorescence

Paraffin embedded specimens were sectioned into 5 µm slices and deparaffinised at 60°C for 2 hours. Simultaneous rehydration and antigen retrieval were achieved with the Antigen Retrieval Buffer (ab93678, Abcam) according to manufacturer’s protocol inside a pressure cooker for 30 minutes. Slides were washed 3 times in PBS for 5 minutes and permeabilised with 0.1% Triton-X in PBS for 10 minutes. Slides were washed and blocked for 30 minutes in 5% goat serum in PBST (PBS+0.1% Tween-20) with 22.52 mg/mL glycine. Samples were stained with mouse anti-CD14 antibody (1:50 dilution) for 1 hour at room temperature and subsequently with mouse anti-CD14 antibody (1:100 dilution) and rabbit anti-PDPN antibody (1:150 dilution) overnight at 4°C. Samples were washed in PBS 3 times for 5 minutes, stained with fluorophore-conjugated secondary antibodies (1:500 dilution, supplementary table S2) and DAPI (1:1000 dilution) for 1 hour at room temperature, washed again in PBS, and mounted with Fluoromount (F4680, Merck). Imaging was accomplished in a Ti2 Eclipse (Nikon) inverted microscope with an iXon Ultra 897 EMCCD camera (Andor).

Images were analysed using Fiji (ImageJ) to quantify mean fluorescence intensity of PDPN expression (mean grey value of the corresponding channel). To quantify cell co-occurrence, immunofluorescence images were processed through a CellProfiler pipeline. Prior to analysis, a mask was manually generated in ImageJ to exclude intestinal crypts. Then, fibroblasts and monocyte segmentation and detection were achieved via thresholding on their responsive channels using a global Otsu method (IdentifyPrimaryObjects). Finally, the mean number of neighbours was quantified using the MeasureObjectNeighbors function with a radius of 200 pixels.

### Statistical Analysis

Individual data points reported in this study represent independent biological replicates, which include different cultures at different passages for CCD-18Cos, hFBLs and THP-1, and different donors for PBMCs and biopsies. Except for the bulk RNAseq data, which only used CCD-18Cos for co-culture, every experiment using fibroblasts was conducted with cells from at least 2 different donors. Data was collected from a minimum of 3-5 independent biological replicates, and analysis was carried out in GraphPad Prism 9. Standard deviation is used throughout. Two-tailed t-test or analysis of variance (ANOVA) with Tukey’s post hoc pairwise comparisons were used to compare between two groups or multiple groups, respectively. Significance was set at P < 0.05 and adjusted for multiple comparisons.

## RESULTS

### TWEAK-stimulated fibroblasts polarise monocytes to an inflammatory phenotype and activate heterologous STAT3 signalling

Previous research from our group has shown that TWEAK-stimulated inflammatory colonic fibroblasts can promote monocyte adhesion and activation, with increased CD14 expression and chemokine production, while TWEAK had no direct effect on monocytes^24^. Building on these observations, we fully characterised the effect of fibroblasts on monocyte activation. THP1 cells were co-cultured with either untreated or TWEAK-treated primary human colonic fibroblasts and analysed via bulk RNAseq. Co-culture with untreated colonic fibroblasts had a profound effect on the transcriptomic profile of THP1s, with 612 upregulated and 604 downregulated genes compared to untreated THP1s. This phenotypic shift was further enhanced in THP1s co-cultured with TWEAK-treated fibroblasts, with over 2000 differentially expressed genes (DEGS, 980 upregulated and 1078 downregulated) compared to naïve THP1s (**Figure 1A**), highlighting the role of the stroma as a regulator of immune responses, and the alterations in this communication that result from inflammatory stimulation in fibroblasts.

**Figure 1.**
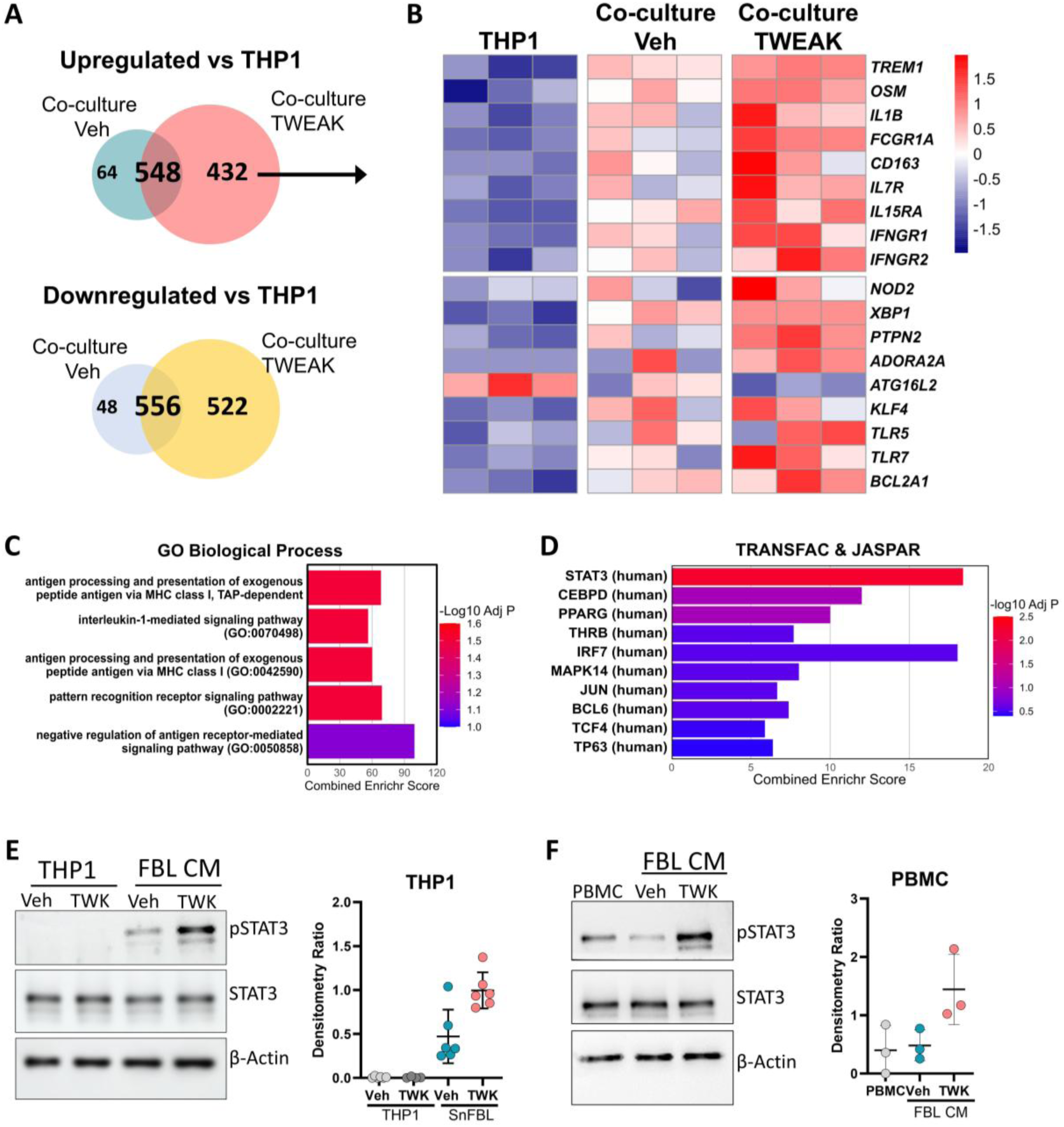
Monocytes polarised in co-culture with TWEAK-activated fibroblasts resemble the phenotype of mononuclear phagocytes in UC. (A) Venn diagrams comparing the DEG upregulated and downregulated in THP1s co-cultured (CC) with Vehicle or TWEAK-treated FBLs compared to naïve THP1s. (B) Heat map of selected genes dysregulated by co-culture with FBLs (n=3, >1.5 fold, p<0.05). (C) (D) Pathway enrichment analysis of transcription factors represented by genes differentially upregulated in THP1s co-cultured with TWEAK-treated fibroblasts. (E-F) Immunoblot analysis and quantification of pSTAT3/STAT3 in THP1s (E) and primary monocytes (F) cultured with conditioned media from human primary colonic fibroblasts (FBL CM) untreated or treated with TWEAK for 48 hours.

We next compared the DEGs upregulated in THP1s co-cultured with healthy vs TWEAK-treated fibroblasts and found an overlap of 548 genes (representing ∼89% of the DEG upregulated by healthy fibroblasts), whereas THP1s co-cultured with TWEAK-treated fibroblasts presented an additional 432 DEGs that were unique (**Figure 1A**). Analysis of this TWEAK-associated signature revealed an increase in the expression of inflammatory macrophage markers (*FCGR1A, CD163*), including genes that have been associated with therapy resistance and fibroblast activation in IBD (*TREM1, OSM, IL1B*) as well as multiple genes associated with susceptibility to IBD (*NOD2, PTPN2, ATG16L1*; **Figure 1B**). Consistent with these findings, pathway enrichment analysis (GO Biological Process) showed an overrepresentation of cellular processes associated with antigen processing and presentation as well as interleukin-1 signalling (**Figure 1C**).

To identify potential mechanisms of fibroblast-monocyte interaction, we performed transcription factor enrichment analysis on the TWEAK-associated gene signature. Cross-referencing with the TRANSFAC and JASPAR databases using the Enrichr tool pointed towards STAT3 as a putative mediator of fibroblast-driven monocyte polarisation (Adjusted p=0.0054; **Figure 1D**). To confirm this, we stimulated both THP1 cells and primary monocytes with conditioned media collected from colonic fibroblasts treated with TWEAK for 48 hours and analysed the phosphorylation levels of STAT3 via immunoblotting. THP1 and primary monocytes stimulated with fibroblast conditioned media showed increased levels of phospho-STAT3 (pSTAT3) compared to naïve cells, which was further increased when the cells were cultured with conditioned media from TWEAK-treated fibroblasts (**Figure 1E-F**). Moreover, differences in STAT3 phosphorylation in THP1s stimulated with conditioned media from TWEAK-treated fibroblasts appeared as early as 1 and 3 hours post-stimulation, pointing to a rapid activation that was sustained overtime (**Supplementary Figure S1**). Together, this data highlights the effects of inflammatory fibroblasts on monocytes differentiation toward an inflammatory profile, which is at least in part mediated by a fibroblast-induced activation of STAT3 signalling in monocytes.

### Blocking non-canonical NF-κB signalling in fibroblasts impairs STAT3 activation and inflammatory response in monocytes

We have previously established that TWEAK can drive the polarisation of colonic fibroblasts at least in part via the non-canonical NF-κB pathway, which can be abrogated through pharmacological inhibition of the NF-κB-inducing kinase (NIK)^24^. Here, we sought to understand whether pharmacological intervention on the fibroblasts could have heterologous effects on monocytes and impair fibroblast-mediated monocyte activation.

Human primary colonic fibroblasts were treated with TWEAK with or without addition of a NIK inhibitor (NIK SMI1) for 48 hours, and their conditioned media was used to stimulate THP1 cells as before (**Figure 2A**). Because we previously found that STAT3 was the top up-regulated pathway in THP1 and confirmed that STAT3 was phosphorylated in both THP1 and primary monocytes in co-culture with TWEAK-treated fibroblasts (**Figure 1E**), we first analysed if NIK SMI1 affected this pathway. Immunoblotting analysis revealed a decrease in STAT3 phosphorylation in THP1s cultured with conditioned media from fibroblasts co-treated with TWEAK and NIK SMI1 compared to those treated with TWEAK alone, down to a level comparable to monocytes treated with conditioned medium from normal fibroblasts (**Figure 2B-C**). We next evaluated the gene expression of inflammatory cytokines (*OSM*, *IL12* and *IL1β*), IBD markers (*NOD2*) and adhesion molecules (*ICAM1*) under the same conditions. Except *ICAM1*, the expression of all targets was significantly increased in THP1 exposed to conditioned media from TWEAK-treated fibroblasts, but not those exposed to conditioned media from naïve fibroblasts, compared to untreated THP1 (**Figure 2D-E**). Concomitant with the loss of STAT3 phosphorylation, THP1 stimulated with conditioned media from fibroblasts co-treated with TWEAK and NIK SMI1 showed a significant decrease in *NOD2* and *OSM* compared to those exposed to conditioned media from TWEAK-treated fibroblasts (**Figure 2D**). Moreover, in the presence of conditioned media from fibroblasts co-treated with TWEAK and NIK SMI1, *IL1β* and *IL12* were not significantly increased compared to naïve THP1, confirming the overall suppressive effect of the NIK SMI1 treatment on the heterologous activation of monocytes (**Figure 2D-E**). The decreased expression of IL-1β was further confirmed at protein level (pro-IL-1β) via immunoblotting (**Figure 2F**). Taken together our data suggest that inhibiting non-canonical NF-κB in inflammatory fibroblasts hijacks their ability to activate STAT3 signalling and inflammatory gene expression in monocytes.

**Figure 2.**
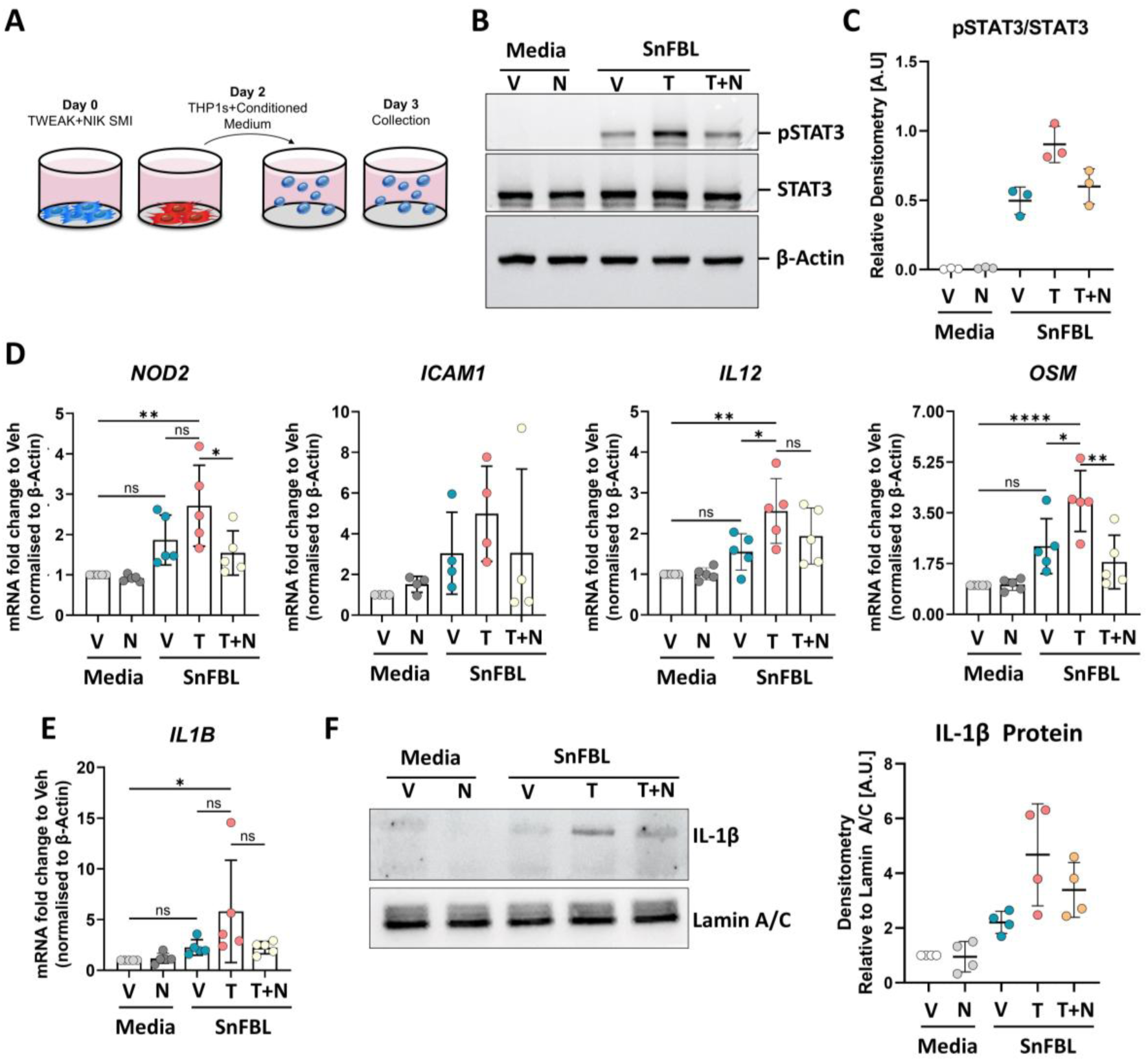
Blocking the non-canonical NF-kB signalling on fibroblasts impairs their ability to activate monocytes. **(A)** Schematic of the indirect co-culture method. Primary colonic fibroblasts were treated with TWEAK alone or TWEAK+NIK inhibitor for 48 hours, and their conditioned medium used to stimulate THP1s for 24 hours. **(B)** Immunoblotting analysis of STAT3 phosphorylation in THP1 cells stimulated with conditioned medium from fibroblasts treated with TWEAK alone or in the presence of the NIK inhibitor NIK SMI01. **(C)** Relative pSTAT3/STAT3 ratio quantified from (B) via densitometry analysis. **(D-E)** Relative expression (mRNA) of *NOD2*, *ICAM1*, *IL12*, *OSM*, and *IL1B* in THP1s stimulated with conditioned medium from fibroblasts treated with TWEAK alone or in the presence of the NIK inhibitor, quantified by qPCR. **(F)** Immunoblotting analysis and quantification of IL1B expression at protein level in THP1s stimulated with conditioned medium from fibroblasts.

### Monocytes polarised by TWEAK-treated fibroblasts *in vitro* resemble the mononuclear phagocyte phenotype in UC

We have previously shown that TWEAK can induce a transcriptional programme in colonic fibroblasts that resembles the inflammatory stroma in UC^17,24^. Given the large number of DEGs we observed in THP1s co-cultured with these fibroblasts, we next sought to investigate whether their transcriptomic profile is representative of monocytes/macrophages in UC. To this end, we decided to explore how the TWEAK-associated THP1 gene signature aligns with the transcriptomic profile of mononuclear phagocyte populations identified in UC patients via single-cell RNA sequencing.

Reanalysis of a scRNAseq atlas of UC biopsies (Ref^18^, Broad DUOS Accession ID: DUOS-000110) identified 15 distinct cell clusters, which could be broadly divided into epithelial, immune, and stromal compartments (**Figure 3A**). Within the myeloid cluster, mononuclear phagocytes (*CSF1R*+) were selected for further downstream analysis. Unbiased clustering of the mononuclear phagocytes identified 11 clusters, including different subpopulations of monocytes, macrophages and dendritic cells (**Figure 3B-D**). Clusters 0 and 1, corresponding to mature or resident macrophages (*C1QC/APOE*) were abundant in both healthy controls and UC biopsies (**Figure 3B-C**), and could be differentiated by the expression of CD209 (Cluster 0). Conversely, Cluster 2 and 3, corresponding to *S100A8/9*+ monocytes expressing high levels of *VCAN* and *FCN1*, was largely absent in healthy controls, but greatly expanded in UC biopsies, consistent with the increased *de novo* infiltrated monocytes. Similarly, Cluster 5 (**Figure 3C**) expressed intermediate levels of both monocyte (*S100A8/9*, *S100A12*) and macrophage (*SOD2*) markers, pointing towards a monocyte-macrophage intermediate or immature macrophage state. Pseudotime analysis of the inflamed UC samples further confirmed Cluster 5 as an early macrophage intermediate and Cluster 0 and 1 as mature macrophage states (**Figure 3E**).

**Figure 3.**
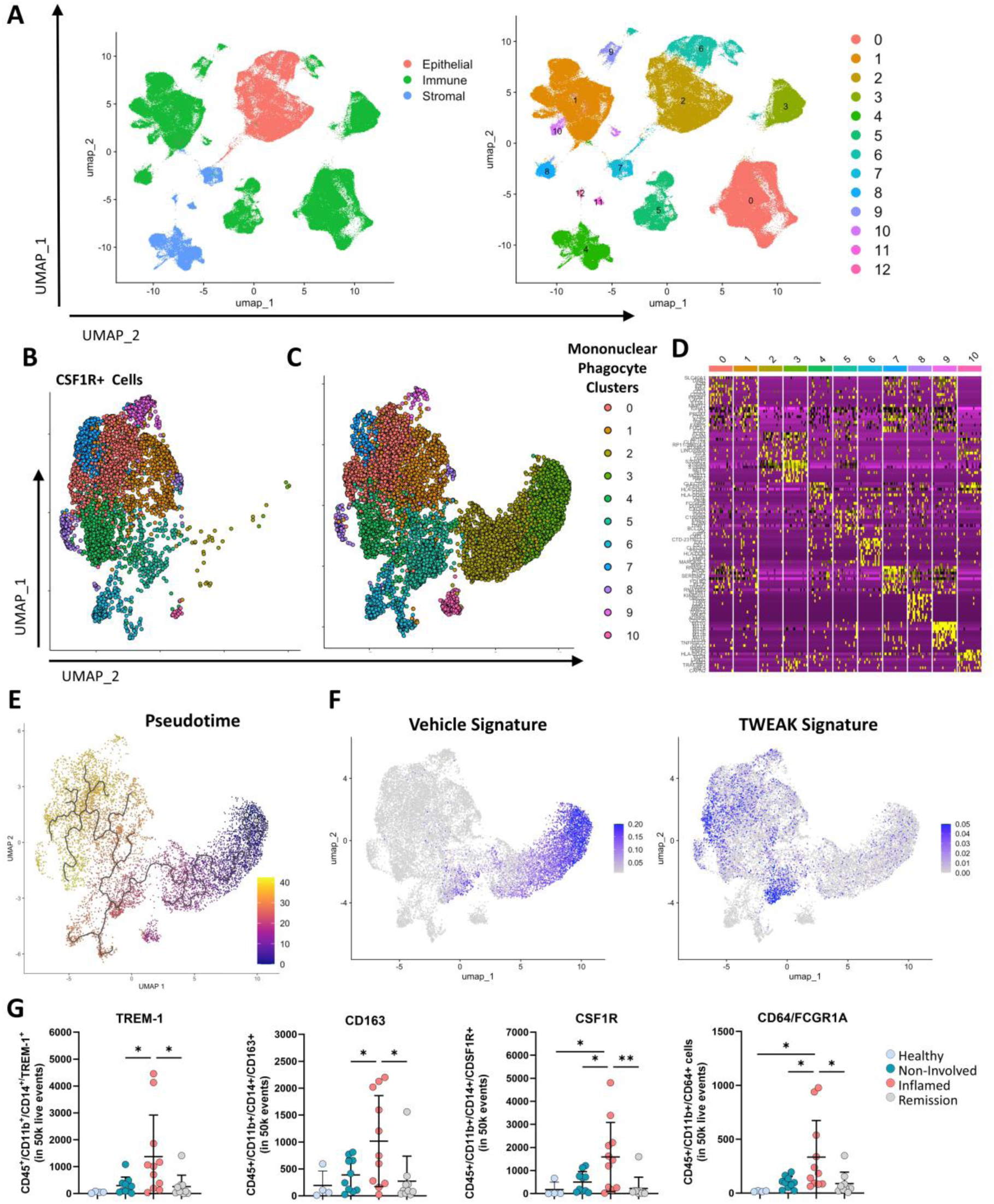
Monocytes polarised in co-culture with TWEAK-activated fibroblasts resemble the phenotype of mononuclear phagocytes in UC. **(A)** UMAP dimensionality reduction analysis of scRNA-seq data from healthy and UC (inflamed and non-involved) biopsies. Reanalysed from Smilie et al. 2019. **(B-D)** UMAP dimensionality reduction analysis of the mononuclear phagocyte (MNP) cluster (CSF1+ cells) identified 11 cell clusters in the colonic mucosal of healthy donors and UC patients. **(E)** Diffusion pseudotime analysis of the MNP clusters in inflamed UC biopsies. **(F)** Expression score for the vehicle and TWEAK-associated modules (defined as the top 50 upregulated genes). **(G)** Frequency of TREM-1^+^, CD163^+,^ CSF1R^+^ and CD64^+^ in mononuclear phagocytes (CD45^hi^/CD11b^+^/CD14^+^ cells) in the colonic mucosa from healthy donors or UC patients with either active disease (inflamed and non-involved sites) or in remission. * p<0.05, ** p<0.01, n≥4.

To investigate the transcriptional alignment between THP1 cells co-cultured with colonic fibroblasts and the mononuclear phagocyte subpopulations identified from the UC scRNAseq atlas, we defined a module score based on the expression of the top 50 most upregulated genes (by fold-change) in THP1s co-cultured with normal fibroblasts (Vehicle Score) or within the TWEAK-associated signature (defined as genes upregulated in THP1s co-cultured with TWEAK-treated fibroblasts but not with normal fibroblasts). Most of the cells aligned with the Vehicle module mapped to Clusters 2 and 3 (infiltrating monocytes), indicating that co-culture with normal colonic fibroblasts may polarise THP1 cells towards a colonic monocyte-like profile (**Figure 3F**). Conversely, cells expressing the TWEAK-module mapped to Clusters 5 (early macrophages) and, to a lesser extent, Cluster 0 (mature CD209 macrophages; **Figure 3F)**. Together, this suggests that co-culture with inflammatory fibroblasts may facilitate the differentiation of monocytes towards a profile that resembles an intermediary state that accumulates in the mucosa of inflamed UC patients.

To validate some of these findings, we used flow cytometry to analyse the mononuclear phagocyte population in mucosal biopsies from UC patients (active or remission) and healthy donors. These cells were gated as CD45^hi^CD14^hi^CD11b^hi^, and expanded in UC active samples compared to matched non-involved samples (**Supplementary Figure S2A).** Consistent with the transcriptomics analysis, we found an accumulation of mononuclear phagocytes expressing TREM1, CD163, CSF1R and CD64 in inflamed biopsies compared with patients in remission or matched non-involved samples (**Figure 3G**). Thus, we find that TWEAK-treated fibroblasts promote monocyte differentiation to a phenotype similar to a macrophage intermediate, and several TWEAK-fibroblast-driven markers are found in infiltrating monocyte/monocyte-derived macrophages in UC.

### TWEAK+ granulocytes accumulate in the colonic mucosa of ulcerative colitis patients

To continue exploring the connection between TWEAK and the inflammatory stroma *ex vivo*, we decided to assess the presence of TWEAK-expressing cells in the mucosa from UC using matched specimens as those in Figure 3G. We first validated the detection of TWEAK by flow cytometry comparing non-stained, FMO control stained and anti-TWEAK stained in CD45^hi^ cells from biopsy specimens (**Figure 4A**). Our flow cytometry analysis showed that the frequency of TWEAK^+^ cells within the leukocyte population (CD45^hi^) increased in inflamed UC biopsies compared to healthy controls, but no significant change in frequency was observed among inflamed, non-involved or remission biopsies (**Figure 4B** and **Supplementary Figure S2B**). Conversely, analysis of the number of TWEAK^+^ myeloid cells (CD45^hi^/CD11b^+^/TWEAK^+^) revealed a significant accumulation in inflamed UC mucosal samples compared to non-involved, remission or healthy control samples (**Figure 4C**). Interestingly, analysis of the same population in CD revealed a lot of variability, with at least 2 patients displaying major increases in the number of CD45^hi^/CD11b^+^/TWEAK^+^ and other 4 patients showing no increase (**Supplementary Figure S3A**). This identifies myeloid cells as a consistent source of TWEAK in UC, but not CD.

**Figure 4.**
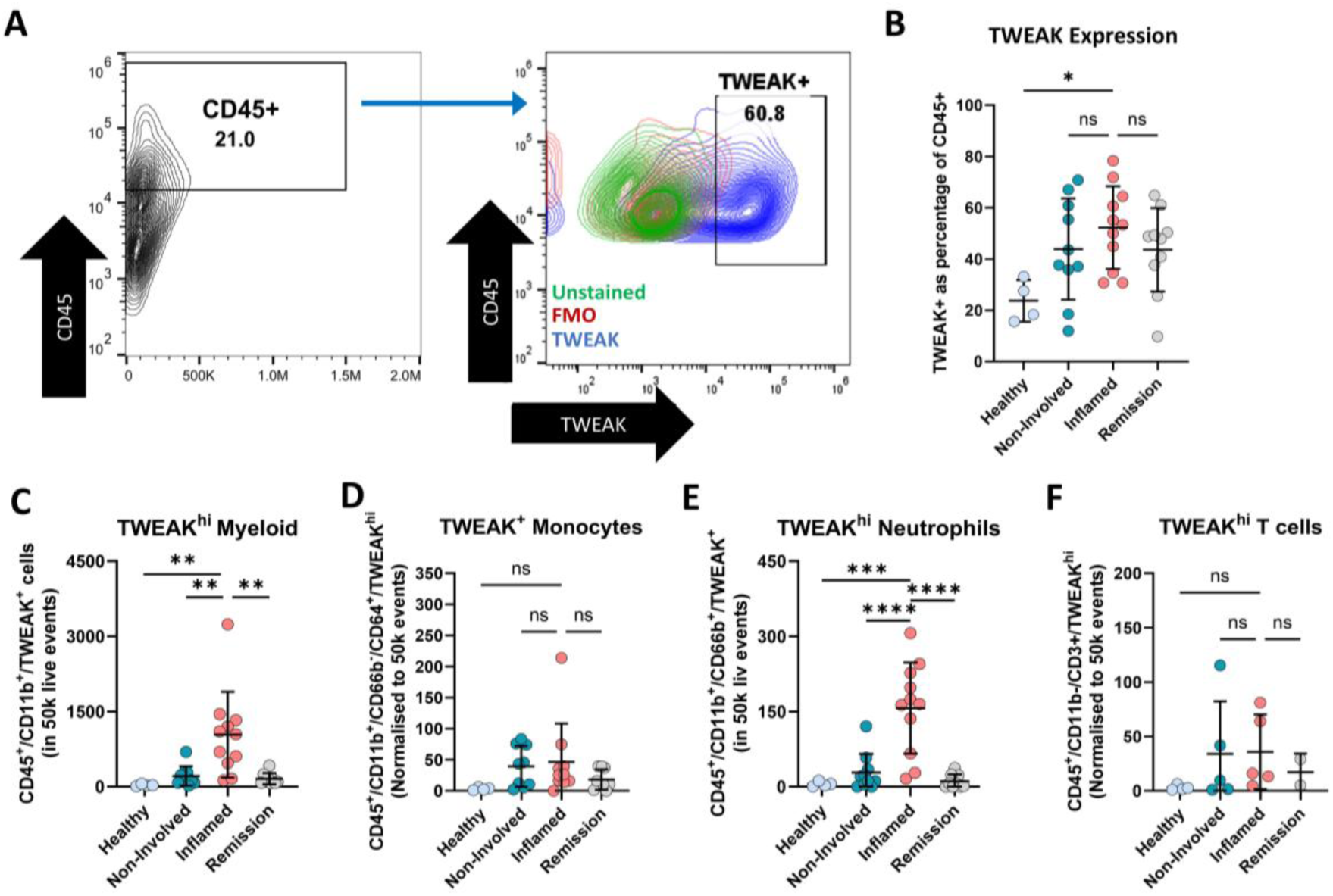
TWEAK+ cells accumulate in UC mucosa. **(A)** Gating strategy used to identify TWEAK^+^ immune cells (CD45^+^), myeloid cells (CD45^+^/CD11b^+^) and granulocytes (CD45^+^/CD11b^+^/CD66b^+^). **(B)** Frequency of TWEAK+ immune cells (relative to the number of CD45^+^ cells) in the colonic mucosa from healthy donors or UC patients with either active disease (inflamed and non-involved sites) or in remission. * p<0.05, n≥4. **(C-F)** Number of TWEAK^+^ myeloid cells (C), monocytes (D), granulocytes (E) and T cells (F) identified in the previous biopsies via flow cytometry. ** p<0.01, *** p<0.001, n≥4.

To understand whether TWEAK was associated with a specific immune cell population in UC, we further characterised myeloid subpopulations, including granulocytes (CD11b^+^/CD66b^+^), monocytes/macrophages (CD11b^+^/CD64^+^/CD66b^-^) and T lymphocytes (CD11b^-^/CD3^+^). While all three populations contained a fraction of TWEAK^+^ cells, we only found significant changes in the abundance of TWEAK^+^ granulocytes (CD45^+^/CD11b^+^/CD66b^+^/TWEAK^+^) when comparing inflamed to matched non-involved UC samples or compared to healthy or remission specimens (**Figure 4D-F**). Our data shows the presence of TWEAK in UC, which may increase in active disease potentially due to the infiltration of TWEAK-expressing granulocytes.

### Fn14+ inflammatory stroma expansion in ulcerative colitis correlates with TWEAK+ cell accumulation and monocyte infiltration

Our previous work showed that TWEAK can polarise colonic fibroblasts *in vitro* towards an inflammatory phenotype characterised by the expression of chemokines and adhesion molecules such as VCAM-1 and PDPN^24^. Given the increased abundance of TWEAK^+^ myeloid cells in the inflamed mucosa of UC patients, we decided to assess the stromal compartment in matched specimens.

Flow cytometry analysis of CD45^-^/EpCAM^-^ cells revealed an increase in the frequency of CD90^+^/PDPN^+^ (**Figure 5A-B, Supplementary Figure S2C**) fibroblasts, consistent with the expansion of the inflammatory stroma previously reported in UC^17,18^. Notably, we also appreciated an increase in these CD90^+^/PDPN^+^ cells in CD specimens (**Supplementary Figure S3B**), consistent with the presence of an inflammatory stromal population reported by others^19,38^. Moreover, when comparing the stromal and myeloid compartments from the same patients, we found a positive correlation between the frequency of CD90^+^/PDPN^+^ cells and the number of TWEAK^+^ myeloid cells (r = 0.7819, p<0.0001; **Figure 5C**). In keeping with this, an analysis of stromal cell compartments in the single cell data reported in Figure 2, showed the accumulation of inflammatory fibroblasts (Cluster 3 fibroblasts), which were characterized by expression of CHI3L1, PTGES, CXCL1, CXCL6, TNFRSF11B and various MMPs (**Figure 5D** and **Supplementary Figure S4**). We then characterized the expression of the TWEAK receptor TNFRSF12A across stromal populations, comparing it to IL1R1 or OSMR which were previously shown to play a role in activating fibroblasts in therapy resistant UC cohorts. TNFRSF12A was highly expressed by numerous cells in the inflammatory cluster 3 (CHI3L1^+^ Fibroblasts), with little to no representation in cells in other stromal clusters (**Figure 5E**). Compared to TNFRSF12A, IL1R1 and OSMR were expressed diffusely and represented in a variety of different fibroblast types, with OSM-R being the least expressed of the 3 receptors (**Figure 5E**). Further analysis also revealed elevated expression of the non-canonical NF-κB components NFKB2 (p100/p52) and RELB along with TNFRSF12A (Fn14) and PDPN in cluster 3 (CHI3L1^+^ Fibroblasts; **Figure 5F**), in agreement with previous findings indicating that TWEAK activates non-canonical NF-kB signalling^24,39^. Given that this population of Fn14+, non-canonical NF-kB-expressing inflammatory fibroblasts emerge in disease, we hypothesised that other classical inflammatory cytokines could promote the expression of Fn14 in naïve fibroblasts. In line with this, we found that the expression of Fn14 in human primary colonic fibroblasts highly increases in response to TNF (**Figure 5G-H**). Taken together our data points towards the role of TWEAK as a driver of inflammatory stroma polarisation in UC.

**Figure 5.**
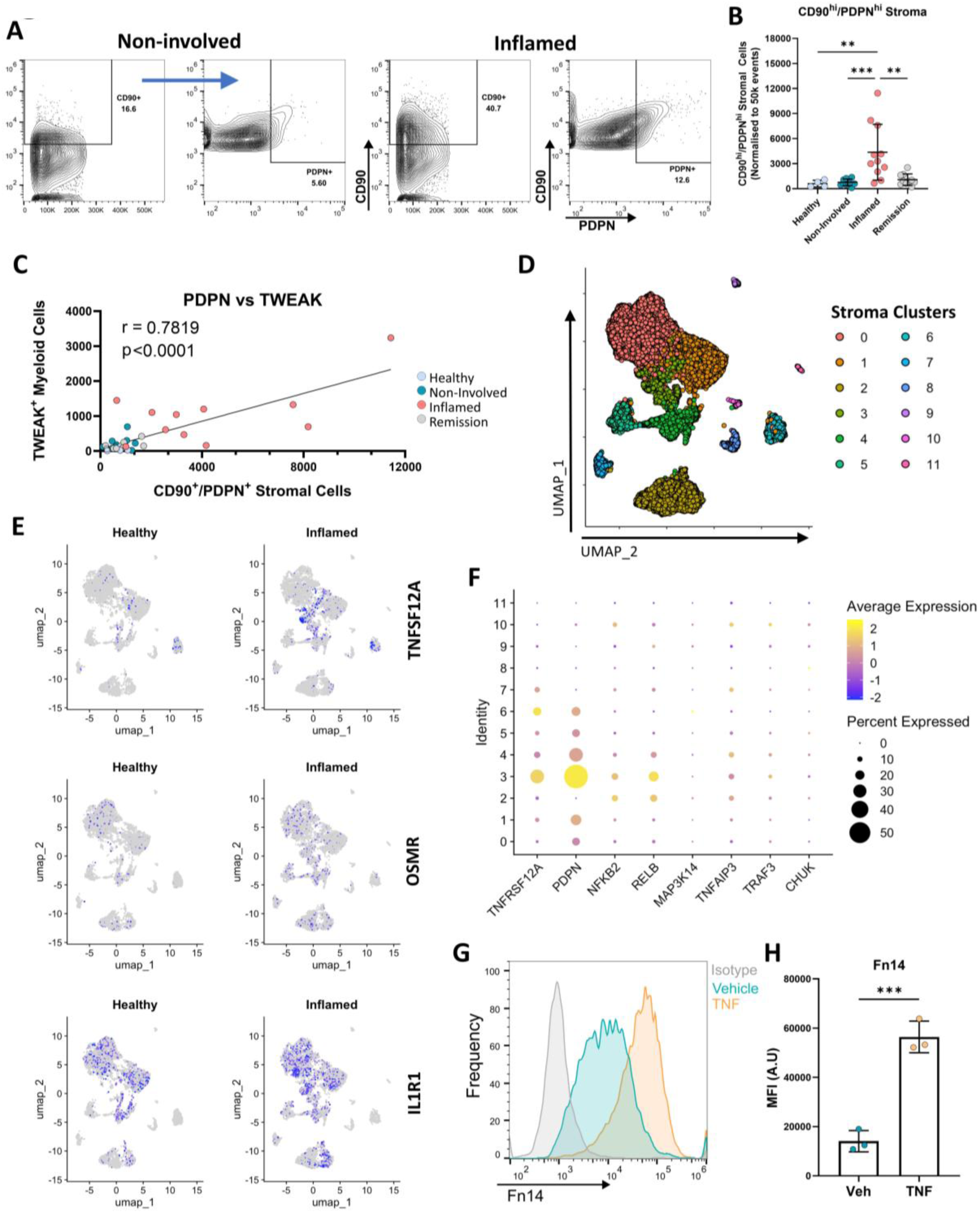
Inflammatory stroma expansion in UC correlates with TWEAK. **(A)** Flow cytometry analysis of CD90^+^/PDPN^+^ stromal cells (CD45^-^/EpCAM^-^) isolated from the colonic mucosa of healthy donors or UC patients with either active disease (inflamed and non-involved sites) or in remission. **(B)** Quantification of the frequency of CD90^+^/PDPN^+^ stromal cells in the previous biopsies (normalised to 50k stromal cells). **(C)** Correlation between the frequency of CD90^+^/PDPN^+^ stromal cells and the abundance of TWEAK^+^ myeloid cells. **(D)** UMAP showing the different stromal cell clusters identified by scRNAseq (reanalysed from *Smillie et al 2019*) **(E)** Expression of Fn14 (TNFRSF12A), OSMR and IL1R1 within the stroma in healthy and inflamed UC mucosa. Reanalysed from Smillie et al 2019**. (F)** Dot plot showing the level (colour) and percentage (size) of expression of the TWEAK receptor (Fn14), Podoplanin (PDPN), and key components of the non-canonical NF-κB signalling pathway. **(G)** Histograms showing the expression of Fn14 by flow cytometry in naive colonic fibroblasts and colonic fibroblasts treated with 5 ng/ml TNF for 24h. **(H)** Quantification of the mean fluorescence intensity (geometric mean) from (F). ** p<0.01, *** p<0.001, **** p<0.0001, n≥4.

Given the ability of inflammatory fibroblasts to promote monocyte adhesion and recruitment, we hypothesised that the expansion of the inflammatory stroma could be accompanied by increased monocyte infiltration and that these cells would co-localize with PDPN^+^ stroma. To further investigate the spatial co-occurrence between these two cell types, we used immunofluorescence (IF) analysis. IF microscopy confirmed the expansion of PDPN^+^ cells in the lamina propria of inflamed UC biopsies compared to matched non-involved controls (**Figure 6A-B**), which was accompanied by an enrichment in CD14^+^ cells located in close proximity to the PDPN^+^ stromal cells (**Figure 6C**). Similarly, based on our flow cytometry data, we found a positive correlation between the frequency of CD90^+^/PDPN^+^ stroma and the number of monocytes (CD45^+^/CD11b^+^/CD14^+^) in the mucosa of UC patients (**Figure 6D**). Taken together, our analysis of IBD specimens points towards the interaction between inflammatory fibroblasts and monocytes, consistent with our *in vitro* data.

**Figure 6.**
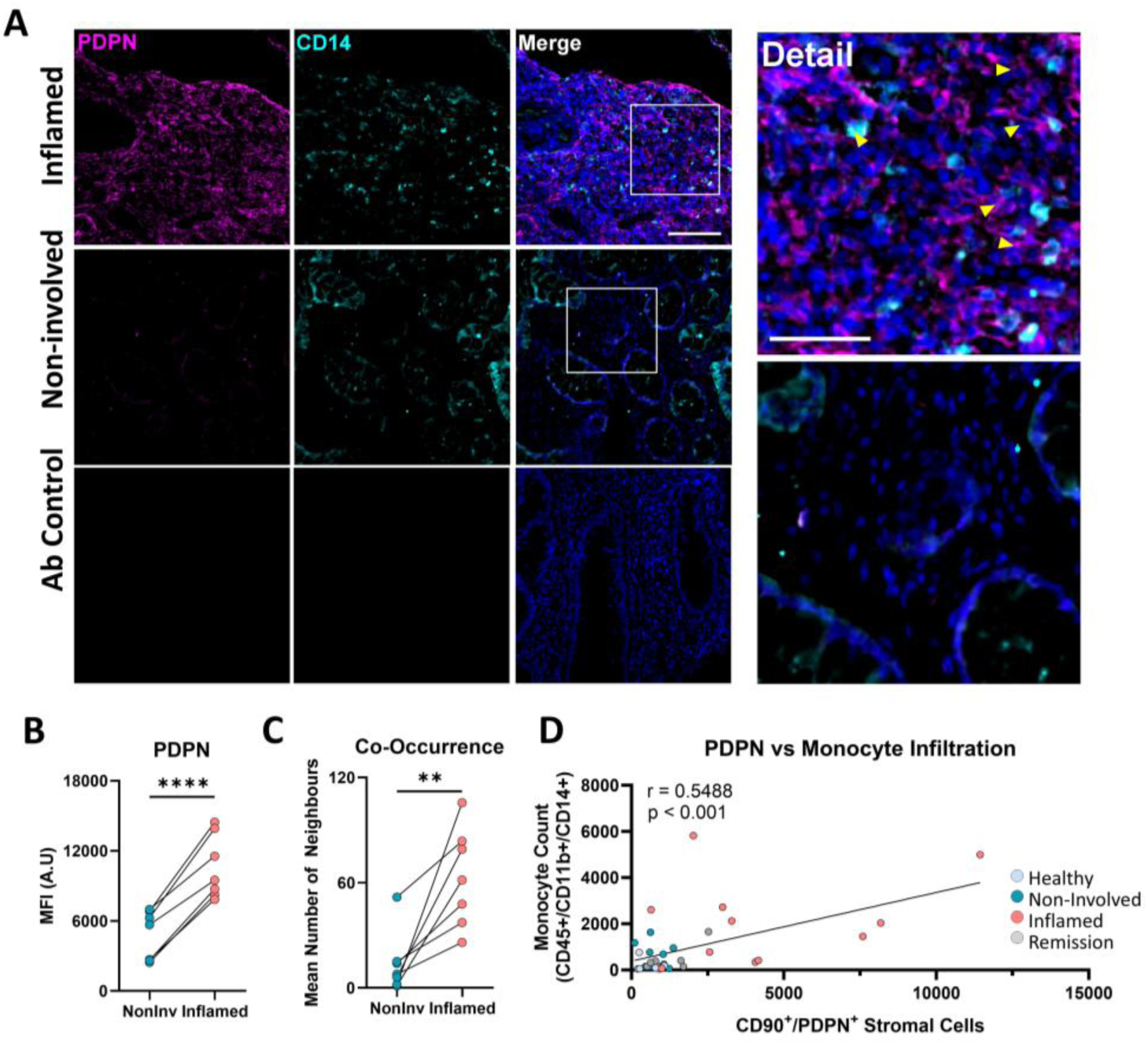
Inflammatory fibroblasts co-localise with infiltrating monocytes in UC mucosa. **(A)** Immunofluorescence staining of inflammatory fibroblasts (PDPN, Magenta) and monocytes (CD14, cyan) in FFPE colonoscopy biopsies from UC patients. Detail panel shows co-occurrence between fibroblasts and monocytes (yellow arrows). **(B)** Quantification of PDPN expression by mean fluorescence intensity from (A). **(C)** Quantification of the average number of monocytes in the vicinity of fibroblasts within a field of view in (A). **(D)** Correlation between the frequency of CD90^+^/PDPN^+^ stromal cells and monocytes (CD45^+^/CD11b^+^/CD14^+^) in biopsies from UC patients and healthy donors. ** p<0.01, n≥5.

## DISCUSSION

Intestinal inflammation in UC is accompanied by a profound remodelling of the cellular composition of the mucosa, including metabolic rewiring of epithelial cells, phenotypic shifts in T and B cells, and the emergence of inflammatory fibroblasts and monocytes^18^. In the complex cellular landscape of UC, the interactions between different inflammation-associated populations have come into the limelight, with several transcriptomics studies identifying fibroblasts-monocyte crosstalk as a central inflammatory hub^18,40^. However, the mechanisms that mediate this cell-cell communication remain incompletely understood. In this manuscript we identify a mechanism of stroma-monocyte communication through a heterologous

TWEAK/NF-kB/STAT3 signalling axis that appears dysregulated in ulcerative colitis, potentially contributing to macrophage dysfunction. Using human endoscopic biopsies, we show that the expansion of CD90^+^/PDPN^+^ inflammatory stroma in the colonic mucosa correlates with the accumulation of TWEAK^+^ myeloid cells, and with the infiltration of monocytes expressing markers relevant in UC such as TREM-1 or CD64. *In vitro*, monocytes co-cultured with TWEAK-stimulated fibroblasts acquired a UC-like phenotype (OSM, IL1B, NOD2) characterized by STAT3 activation, which can be depleted via selective inhibition of non-canonical NF-κB pathway on the fibroblasts.

Since the identification of inflammatory fibroblasts in IBD, multiple cytokine/receptor pairs have been accredited with contributing to their polarisation. West *et al*. found that the member of the IL-6 family oncostatin M (OSM) induces the expression of cytokines and cell adhesion molecules in colonic fibroblasts, and that its receptor OSM-R is highly upregulated in IBD stroma^20^. The same group later identified the IL-1β/IL-1R1 pair as a potent activator of fibroblasts in IBD, in turn promoting neutrophil recruitment^16^. Interestingly, both IL-1β/IL-1R1 and OSM/OSM-R expression correlated with disease severity and resistance to anti-TNF, suggesting that stroma-immune interaction may play a crucial role in the response to therapy. More recently Cadinu *et al.* identified several putative cytokines that could contribute to the interaction between inflammatory and normal fibroblast in a DSS-induced murine colitis, including *il11*, *lif*, or *tnfsf13b* (BAFF), suggesting that inflammatory polarisation could propagate through the stroma^41^. One significant limitation in our understanding of stromal remodelling in IBD is the use of endoscopic biopsies. More recently, analysis of transmural samples from CD patients revealed a population of CD-specific FAP^+^ fibroblasts involved in fibrostenosis^42^. This population of activated fibroblasts closely co-localised with, and was activated by, CD150^+^ inflammatory monocytes via TWIST1. Our in vitro co-culture model, while limited to an *in vitro* context, closely recapitulates the transcriptional programmes of inflammatory fibroblasts and monocytes in UC, and cloud therefore constitute a powerful platform to dissect the complex and bi-directional stroma-immune crosstalk in IBD.

Our work positions TWEAK and its cognate receptor Fn14 as another contributor to inflammatory stroma polarisation in UC, promoting fibroblast-mediated monocyte differentiation towards a phenotype that resembles early monocyte/macrophage intermediates. Interestingly, based on scRNA-seq data, the expression of the receptor Fn14 seems relatively specific to fibroblast populations that emerge in UC, compared to other receptors such as OSM-R and IL-1R1. This is remarkable considering that IL-1, OSM and their receptors have previously been linked to stromal activation and warrants further studies to clarify the full extent to which TWEAK contributes to inflammatory fibroblast function in UC. It is also noteworthy that treatment of naïve primary colonic fibroblast with the IBD-relevant cytokine TNF increased the expression of Fn14. Considering that Fn14 displayed relatively marginal expression in healthy tissues, but was abundant in stromal cells in UC, it is tempting to speculate that a TWEAK-Fn14 axis could be enabled as a result of inflammatory activation of fibroblasts, although this will need to be further analysed.

The source of TWEAK in the gut has remained unclear, with its transcript being detected in both immune cells and enterocytes^25^. Our findings here point towards myeloid cells as an important source of TWEAK, and highlight the contribution of granulocytes, which can be underrepresented in transcriptomics data. It is worth to caveat, however, that granulocytes tend to secrete low concentrations of cytokines compared to other myeloid or lymphoid cells. Nevertheless, TWEAK has also been found as a membrane bound product, which was shown to activate signalling in neighbouring cells^43^. Since granulocytes, and particularly neutrophils, are abundantly recruited to the UC mucosa and have been found to interact with fibroblasts, these interactions could allow for a neutrophil-mediated activation of TWEAK signalling in fibroblasts^16^. Moreover, our data also showed a high degree of variability in the levels of

TWEAK in CD patients. Whether this is the result of the diversity of CD endotypes, tissue locations or therapy status, and if TWEAK could contribute to CD in a specific tissue region or disease stage remains to be determined. However, in the emerging and ever-growing list of stroma-relevant cytokines, it is unlikely that a single mediator is responsible for the activation of inflammatory fibroblasts. Thus, understanding which cytokines dominate the stromal immune response, and whether a hierarchy exists for these mediators in different disease contexts, could constitute a step towards patient stratification in UC.

We and others have previously reported that inflammatory fibroblasts can recruit and activate monocytes *in vitro*^21,24^, but the mechanisms have not been characterised. Here, we found inflammatory fibroblast-dependent activation of monocytes is driven at least in part by STAT3, consistent with another report that detected increased baseline levels of pSTAT3 in monocytes from UC patients compared to CD or healthy controls^44^. The JAK/STAT pathway has garnered increasing interest in the field of IBD owing to their diverse functions in adaptive and innate immunity. Numerous cytokines associated with gut homeostasis (IL-10, IL-22) and IBD pathogenesis (IL-6, IL-12, IL-23, GM-CSF) signal through JAK/STATs^45^, and gain of function mutations (e.g. JAK2) are associated with increased susceptibility to IBD^46^. For these reasons, JAK/STATs has become a therapeutic priority in IBD, with the pan-JAK inhibitor Tofacitinib currently approved for UC and several other small molecule inhibitors being in the pipeline. However, while STAT3 constitutes an interesting target, its dual role in the signalling pathways of both pro- and anti-inflammatory cytokines limits its effectiveness. Indeed, analysis of patients refractory to Tofacitinib points towards the impairment of the IL-10/STAT3 pathway in macrophages as a likely culprit^47^, suggesting that direct blockage of STAT3 signalling may be detrimental in some IBD patients.

Given the role of inflammatory fibroblasts in orchestrating monocyte recruitment and activation, we hypothesised that addressing fibroblasts inflammatory polarisation could restore healthy stroma/immune crosstalk. Based on this idea, we found that fibroblasts-dependent STAT3 activation on monocytes can be targeted indirectly by impairing non-canonical NF-κB signalling on inflammatory fibroblasts, which could offer an alternative approach to the direct inhibition of STAT3. Moreover, treatment with an inhibitor against NIK abrogated the upregulation of IL1B, NOD and OSM on monocytes, suggesting that this therapeutic strategy could effectively reprogram fibroblast/monocyte interactions. Interestingly, high levels of non-canonical NF-κB activation have also been linked with therapy resistance in UC patients^48^, and our analysis of sc-RNAseq data points towards the co-expression of non-canonical NF-κB components along with FN14 and PDPN in inflammatory fibroblasts.

Taken together, our data solidifies TWEAK as a driver of inflammatory stroma polarisation and stroma/monocyte crosstalk in UC. Using human biopsies, *in vitro* co-culture models and transcriptional alignment studies, we show that CD90^+^/PDPN^+^ fibroblasts expand in the colonic mucosa of UC patients, co-localise with infiltrating monocytes *in vivo*, and promote monocyte activation and STAT3 phosphorylation *in vitro*. Collectively, our manuscript identifies the TWEAK/NF-κB/STAT3 axis as a novel signalling hub in stroma-monocyte communication, which can be manipulated by targeting non-canonical NF-κB in the stroma.

## Acknowledgements

This work was supported by funding from Science Foundation Ireland and The Irish Research Council under the Pathway program (21/PATH-S/9621), funding from the UCD Ad Astra fellows program and UCD SBBS SPARK awards. CM was supported by the IRC GOI scheme under award number GOIPD/2023/1118. SB was supported by the College of Science STEM summer research scholarship program. We are thankful to all the blood donors. We would like to thank the genomics, flow cytometry and histology CORE facilities at UCD Conway Institute for access and support.

## Data Availability

Raw data from Bulk RNAseq can be accessed at GSE308641.

## Author contributions

C.M and M.C.M drafted this manuscript. C.M and M.C.M designed the study. C.M, B.R, S.B, C.M.O, C.B, and M.B.NÍ C, performed experiments. C.M, G.R.J and C.K analysed sequencing data. M.N, K.D and D.W collected clinical samples. C.A, H.R, C.B, G.D, S.T. and M.C.M have made intellectual and resource contributions. All authors approved the final draft.

