## Supplementary Material for "TWEAK is increased in ulcerative colitis and contributes to fibroblast-mediated monocyte activation via heterologous non-canonical NF-kB/STAT3 signalling"

**Supplementary Table S1. Clinical Data**

| | UC Active | UC Remission | CD Active | Non-IBD |
|---|---|---|---|---|
| Number of Patients | 15 | 10 | 7 | 4 |
| **Age** | | | | |
| Median | 40 | 54 | 26 | 52.5 |
| Range | 26-65 | 20-71 | 17-66 | 28-67 |
| **Sex** | | | | |
| Male | 53% | 60% | 57% | 25% |
| Female | 47% | 40% | 42% | 75% |
| **Severity** | | | | |
| Mayo 1 | 6 | - | - | - |
| Mayo 2 | 6 | - | - | - |
| Mayo 3 | 3 | - | - | - |
| **Treatment** | | | | |
| Anti-TNF | 5 | 3 | 5 | 0 |
| 5ASA | 4 | 2 | 0 | 0 |
| Anti-TNF+5ASA | 1 | 0 | 0 | 0 |
| Jak-i | 2 | 1 | 0 | 0 |
| IMURAN | 1 | 0 | 0 | 0 |
| Vedolizumab | 0 | 1 | 0 | 0 |
| NIL | 2 | 3 | 2 | 4 |

**Supplementary Table S2. Reagent List**

**qPCR Primers**

| Gene | Primer | Sequence 5’–3’ |
|---|---|---|
| **ACTB** | Fwd | CGACAGGATGCAGAAGGAGA |
| | Rev | CATCTGCTGGAAGGTGGACA |
| **NOD2** | Fwd | CTCCGAGGCAACACCTCCTT |
| | Rev | TGCCAATGTTCCCCACC |
| **ICAM1** | Fwd | TTGTTGGGCATAGAGACCCC |
| | Rev | GGTTTTAGCTGTTGACTGCCC |
| **IL12** | Fwd | CCAGGTGGTTCAAGACCAT |
| | Rev | ATCCGGTTCTTTCAAGGGAGG |
| **OSM** | Fwd | AGAGTACCGCGTGCTCCTTG |
| | Rev | CCTGCAGTGCTCTCTCAGTTT |
| **IL1B** | Fwd | CCACTACAGCAAGGGCTTCA |
| | Rev | ATCGTGCACATAAGCCTCGT |

**Immunoblot Antibodies**

| Target | Clone | Supplier | Cat. Number |
|---|---|---|---|
| **β-Actin** | AC-15 | Sigma | A5441 |
| **pSTAT3 (Tyr705)** | D3A7 | Cells Signalling Technologies | 9145 |
| **STAT3** | 124H6 | Cells Signalling Technologies | 9139 |
| **IL-1β** | D3U3E | Cells Signalling Technologies | 12703 |

| | | | |
|---|---|---|---|
| **Lamin A/C** | 4C11 | Cells Signalling Technologies | 4777 |
| **Anti-rabbit HRP** | - | Cell Signalling Technologies | 7074 |
| **Anti-mouse HRP** | - | Cell Signalling Technologies | 7076 |

**Flow Cytometry Antibodies**

| Antibody | Clone | Fluorochrome | Supplier | Cat. No. |
|---|---|---|---|---|
| CD90 | 5E10 | Alexa Fluor 700 | Biolegend | 328119 |
| PDPN | NC-08 | APC/Cyanine7 | Biolegend | 337029 |
| CD45 | 2D1 | APC/Cyanine7 | Biolegend | 368515 |
| EpCAM | 9C4 | APC | Biolegend | 324207 |
| CD45 | HI30 | Pacific Blue | Biolegend | 304021 |
| CD11b | QA20A58 | APC | Biolegend | 379905 |
| CD14 | 63D3 | FITC | Biolegend | 367116 |
| CSF1R | 9-4D2-1E4 | PerCP Cy5.5 | Biolegend | 347309 |
| CD163 | GHI/61 | Bv605 | Biolegend | 333615 |
| TREM-1 | TREM-37 | PE | Biolegend | 316103 |
| CD3 | OKT3 | FITC | Biolegend | 317305 |
| CD66b | G10F5 | PE/Cy7 | Biolegend | 305115 |
| CD64 | 10.1 | PerCP Cy5.5 | Biolegend | 305023 |
| TWEAK | CARL-1 | PE | Biolegend | 308305 |

**Immunofluorescence Antibodies**

| Target | Clone | Supplier | Cat. Number |
|---|---|---|---|
| **Podoplanin** | Polyclonal | Proteintech | 22099-1-AP |
| **CD14** | 2C1D9 | Proteintech | 60253-1-IG |
| **Anti-Rabbit AF568** | - | Thermo Fisher | A10042 |
| **Anti-Mouse AF647** | - | Thermo Fisher | A21236 |

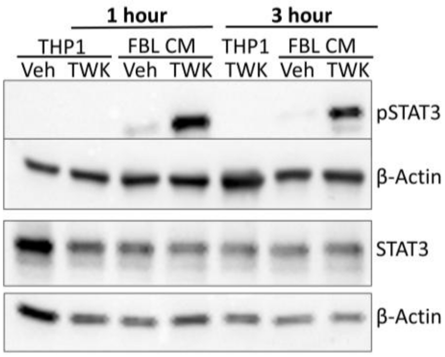


**Supplementary Figure S1. Fibroblasts activate STAT3 signalling in monocytes at early time points.** Immunoblotting analysis of STAT3 phosphorylation in THP1 cells stimulated for 1 hour or 3 hours with either TWEAK directly or conditioned medium from fibroblasts untreated or treated with 50 ng/mL TWEAK for 48h.

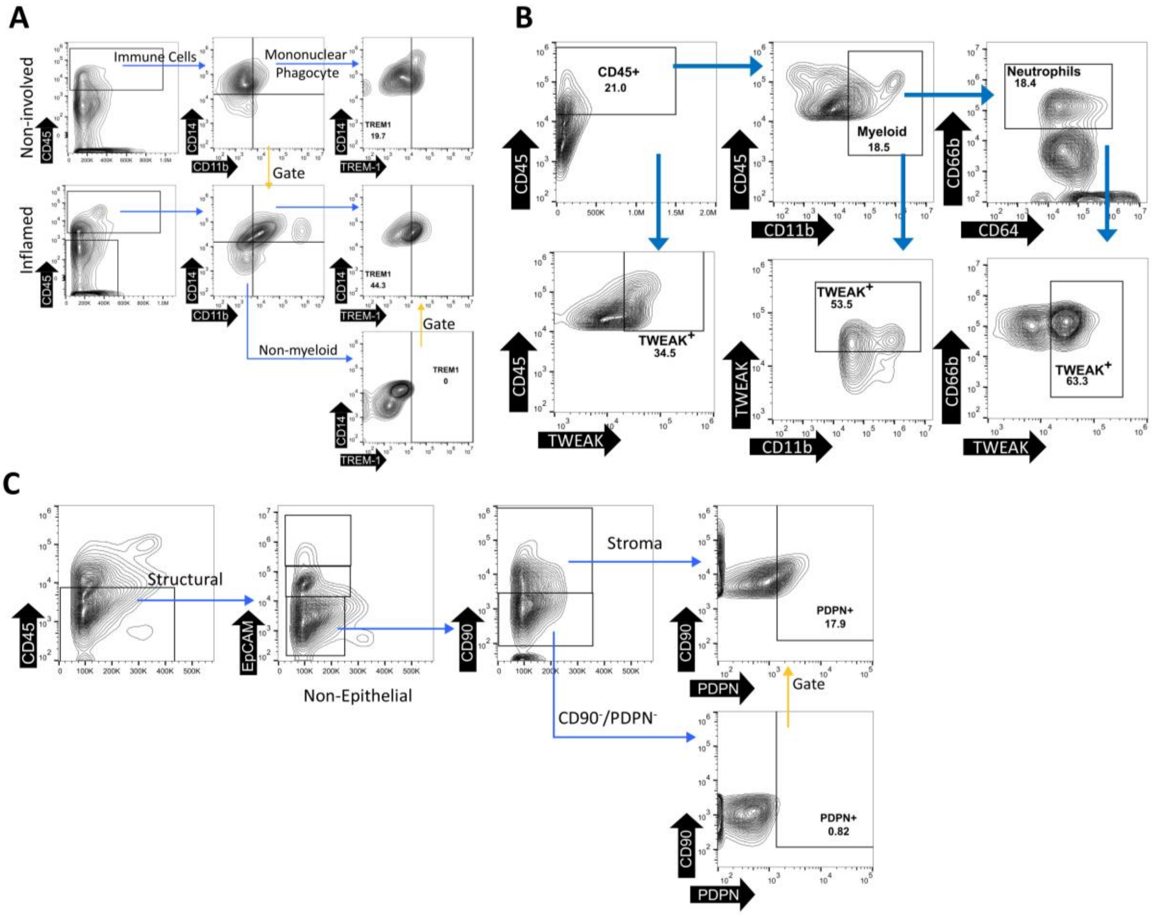


**Supplementary Figure S2. Gating strategy. (A)** Gating strategy used to analyse the mononuclear phagocyte compartments using flow cytometry. **(B)** Gating strategy to identify

TWEAK$^{+}$ circulating granulocytes from whole blood. **(C)** Gating strategy used to analyse the stroma compartment using flow cytometry.

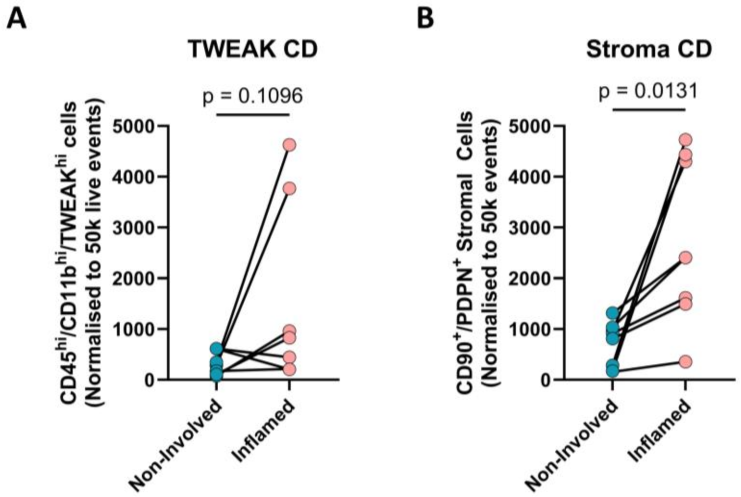


**Supplementary Figure S3. Inflammatory stroma and TWEAK$^{+}$ myeloid cells in CD biopsies. (A)** Number of TWEAK$^{+}$ myeloid cells identified in the previous biopsies. **(B)** Quantification of the frequency of CD90$^{+}$/PDPN$^{+}$ stromal cells in CD biopsies and matched non-involved controls.

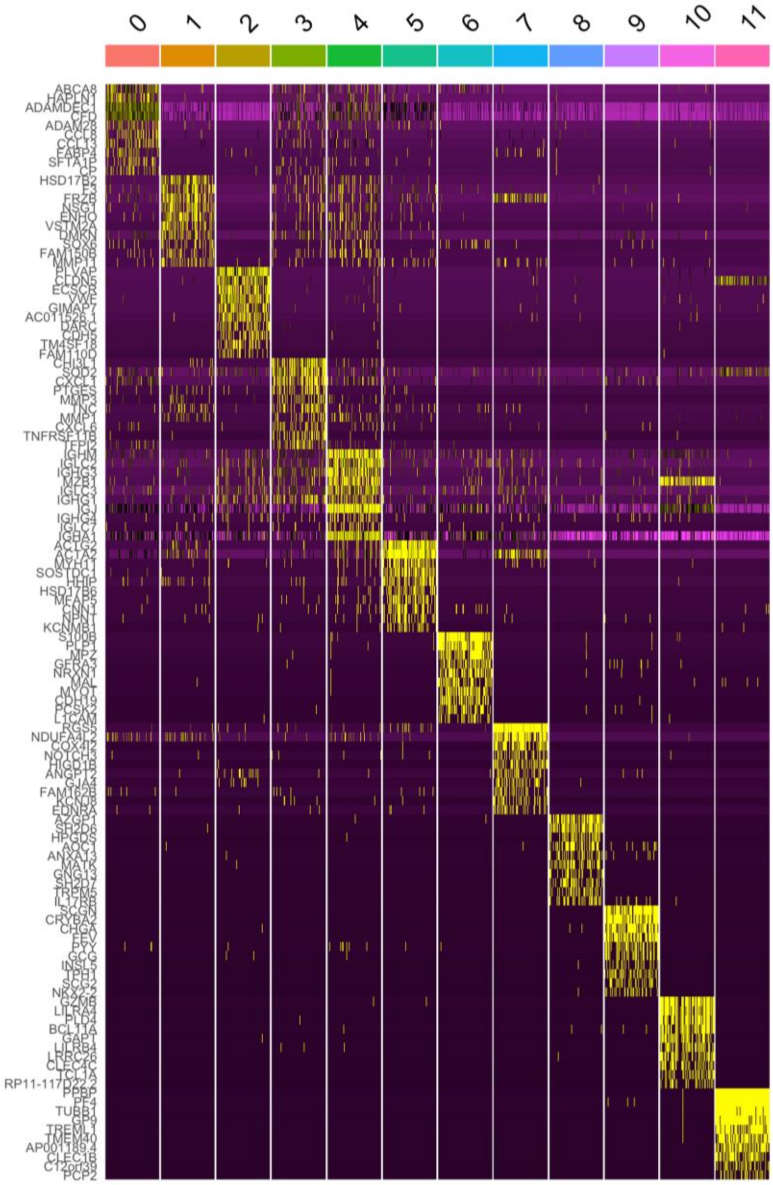


**Supplementary Figure S4.** Definition of stromal clusters in figure 5.
